# Cryo-EM structure of the Hedgehog release protein Dispatched

**DOI:** 10.1101/707513

**Authors:** Fabien Cannac, Chao Qi, Julia Falschlunger, George Hausmann, Konrad Basler, Volodymyr M. Korkhov

## Abstract

The Hedgehog signaling pathway controls embryonic development and adult tissue homeostasis in multicellular organisms. In Drosophila melanogaster, the pathway is primed by secretion of a dually lipid-modified morphogen, Hedgehog (Hh), a process dependent on a membrane-integral protein Dispatched. Although Dispatched is a critical component of the pathway, the structural basis of its activity has so far not been described. Here, we describe a cryo-EM structure of the Drosophila melanogaster Dispatched at 3.2 Å resolution. The ectodomains of Dispatched adopt an open conformation suggestive of a receptor-chaperone role. A 3D reconstruction of Dispatched bound to Hh confirms the ability of Dispatched to bind Hh but using a unique mode distinct from those previously observed in structures of Hh complexes. The structure may represent the state of the complex that precedes shedding of Hh from the surface of the morphogen-releasing cell.

## Introduction

The Hedgehog (Hh) signaling pathway controls embryonic development and adult tissue homeostasis in multicellular organisms. The Hh pathway is a key homeostatic regulator in embryogenic development, tissue regeneration and stem cell maintenance (*1*, *2*). Disruption of this pathway leads to deregulation of tissue homeostasis, developmental diseases and cancer (*3*, *4*). The key players in the Hh pathway are highly conserved across species (*5*), which has led the Drosophila melanogaster Hh system to being one of the main sources of insights into the fundamental mechanisms of Hh signaling regulation. The *D. melanogaster* Hh is synthesized as a pre-protein that is autoproteolytically processed by its C-terminal domain and dually lipidated with an N-terminal palmitoyl and a C-terminal cholesterol moieties (the processed form of Hh, HhN-P, is here referred to as HhN) (*6*-*8*). The lipid modifications presumably anchor the Hh ligand to the biological membranes and extracellular matrix, ensuring that its short- and long-range actions are tightly controlled (*9*, *10*). Binding of HhN to Patched on the surface of the Hedgehog-responding cells alleviates the inhibitory action of Patched on Smoothened, a G protein-coupled receptor-like membrane protein (*11*). This leads to phosphorylation of the Smoothened C-terminus and recruitment of Costal 2 (Cos2) and Fused (Fu), with subsequent activation of the Cubitus interruptus (Ci) transcription factor that induces the transcription of the Hh target genes (*11*).

The Hh pathway is initiated by the release of the HhN from the producing cell (Fig. 1A), and this process is facilitated by vertebrate and invertebrate Dispatched homologues via an undefined mechanism. *Drosophila* Dispatched is a 139 kDa protein predicted to contain twelve transmembrane (TM) helices and two extracellular domains (ECDs) (*12*). Like Patched, Dispatched belongs to the RND family of membrane transporters. The TM2-6 region of Dispatched is annotated as the sterol sensing domain (SSD) (*13*), similar to Patched and other eukaryotic RND proteins. The functional significance of the Dispatched SSD for interaction with HhN is unclear (*14*). The vertebrate Dispatched homologue has been proposed to synergize with Scube2 in facilitating the release of the cholesterol-modified sonic Hedgehog (Shh) (*15*, *16*). Early experiments showed that a possible transporter-like activity of Dispatched may be necessary for its role in Hh release(*17*). Furthermore, a recent report indicated that Shh release by mouse DISP requires processing of the latter by a convertase Furin (*18*). A similar proteolytic activation was proposed for the *D. melanogaster* Dispatched (*18*). Recent structural work on mammalian RND transporters, such as PTCH1 and NPC1, revealed the architecture of these proteins and provided snapshots of these proteins in various states (*19*-*23*). Despite these recent breakthroughs, the precise role of Dispatched in Hedgehog release has remained unclear.

**Fig. 1.**
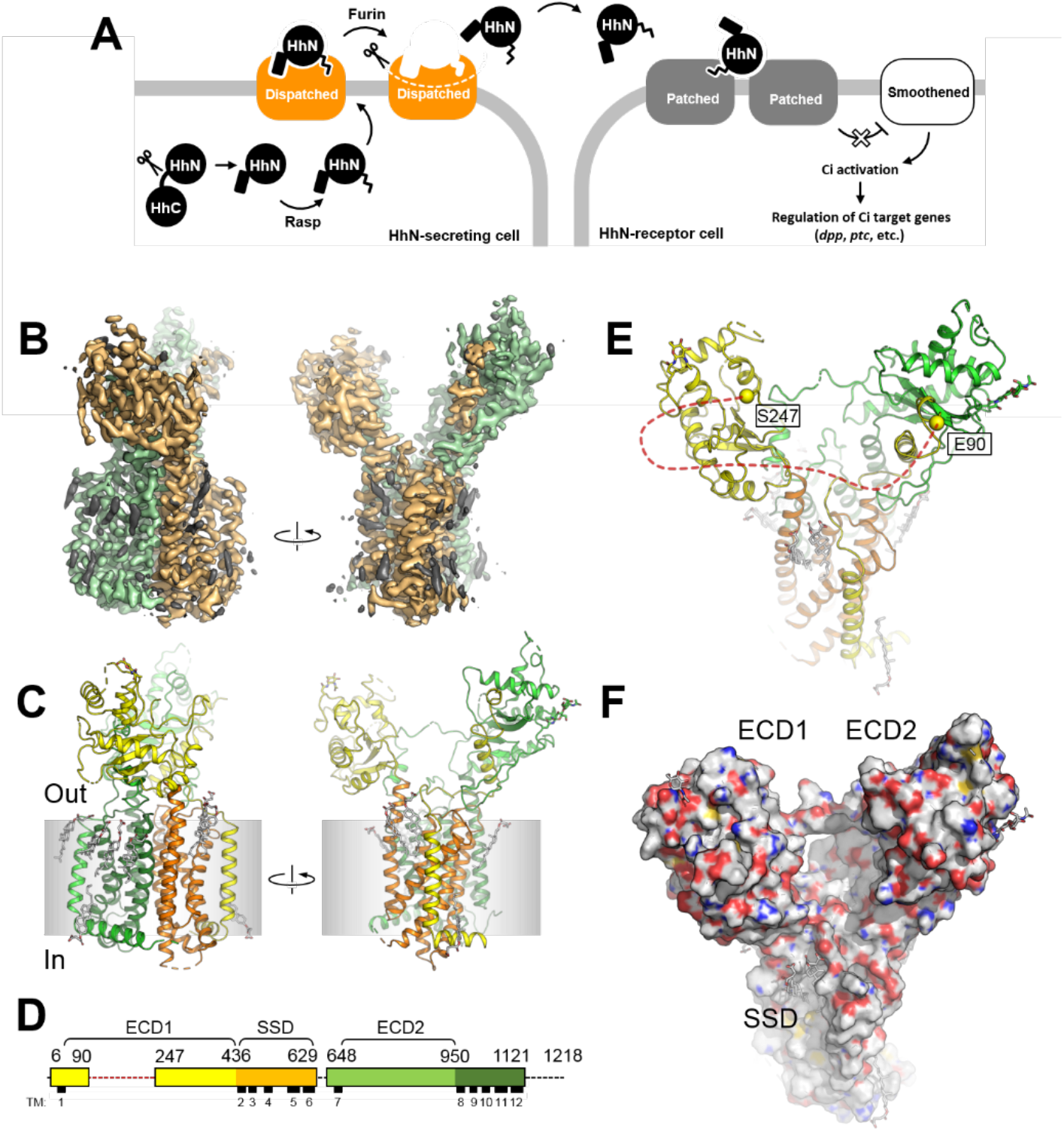
Structure of the *Drosophila melanogaster* Dispatched. (**A**) The schematic representation of Hh ligand release, which initiates the Hh pathway. (**B**) A cryo-EM map of Dispatched reconstructed at 3.2 Å resolution; orange and green colors indicate the N- and C-terminal halves of the protein; elements of the density map assigned to bound sterols or unassigned (i.e. corresponding to the unstructured components of the detergent micelle) are coloured gray. (**C**) The atomic model of Dispatched, built using the 3.2 Å map shown in B. (**D**) Sequence coverage of the atomic model and annotation of the sequence. The colours of the linear representation of Dispatched sequence in D correspond to the colours of the secondary structure elements in c. ECD1 – ectodomain 1; ECD2 – ectodomain 2; SSD – sterol-sensing domain. Dotted line indicates a portion of the protein not resolved in the cryo-EM map. (**E**) The view of the protein model from the extracellular side indicates the Cα atoms of the residues E90 and S247; the red dotted line indicates an unresolved loop between these two residues. (**F**) The space-filling representation of the Dispatched model reveals a large cavity formed by the two ectodomains, with the SSD in close proximity.

We now report cryo-EM structures of the Drosophila melanogaster Dispatched in the apo state and in presence of the Hedgehog ligand. The ectodomain (ECD) of Dispatched splays open in a new conformation. Furthermore, the structure of the complex highlights a new interface of binding for the morphogen. The structure likely represents the state of the complex that precedes shedding of Hh from the surface of the morphogen-releasing cell. The ability of the Dispatched ECD to form a bowl-shaped cavity may indicate that homologous proteins of the Resistance-Nodulation-Division (RND) family of membrane transporters, such as Patched (PTCH1) or Nieman Pick Desease protein 1 (NPC1) may adopt similar conformations.

## Results

### Cryo-EM structure of Dispatched

To gain insight into the structure and function of Dispatched, we expressed the Drosophila Dispatched homologue in HEK293F cells (Fig. S1A), purified the protein by affinity chromatography, and collected a high resolution single particle cryo-EM dataset (Fig. S2A). The excellent quality of the sample allowed us to reconstruct the cryo-EM density map of Dispatched at a resolution of 3.2 Å (Fig. 1B, Fig. S2B-D). Using the high resolution density map of Dispatched, we built the complete model of the protein, including the TM domain bundle and the two ectodomains, covering the majority (~69%) of the Dispatched amino acid sequence (Fig. 1C-D).

The structure revealed the expected TM domain arrangement characteristic of the RND transporters (Fig 1C-D, Fig 2). The density map of Dispatched showed a number of density elements that we interpreted as molecules of bound cholesteryl hemisuccinate (CHS), which was present during protein purification (Fig. 1B, Fig. S3). The SSD of Dispatched seems to have the ability to accommodate several sterol molecules within its pocket region, similar to the SSD of PTCH1 bound to ShhN_C24II_ (*24*). We have modelled a total of 9 sterol molecules bound to the inner and outer leaflet regions at the protein-lipid bilayer interface of Dispatched (Fig. S2). Our reconstruction also revealed a striking feature of the extracellular portion of the protein: the ectodomains of the Dispatched splayed apart creating a large bowl-shaped cavity (Fig. 1E-F). The loop between the residues E90 and S247, which belongs to ECD1, is not resolved in our reconstruction (Fig. 1E). This loop is predicted to be mostly unstructured, which may explain the lack of interpretable cryo-EM density corresponding to this region of the protein. The absence of a connecting protein density between the two ECDs creates an opening into the Dispatched ECD cavity in close proximity to the protein’s membrane-embedded SSD region (Fig. 1F).

**Fig. 2.**
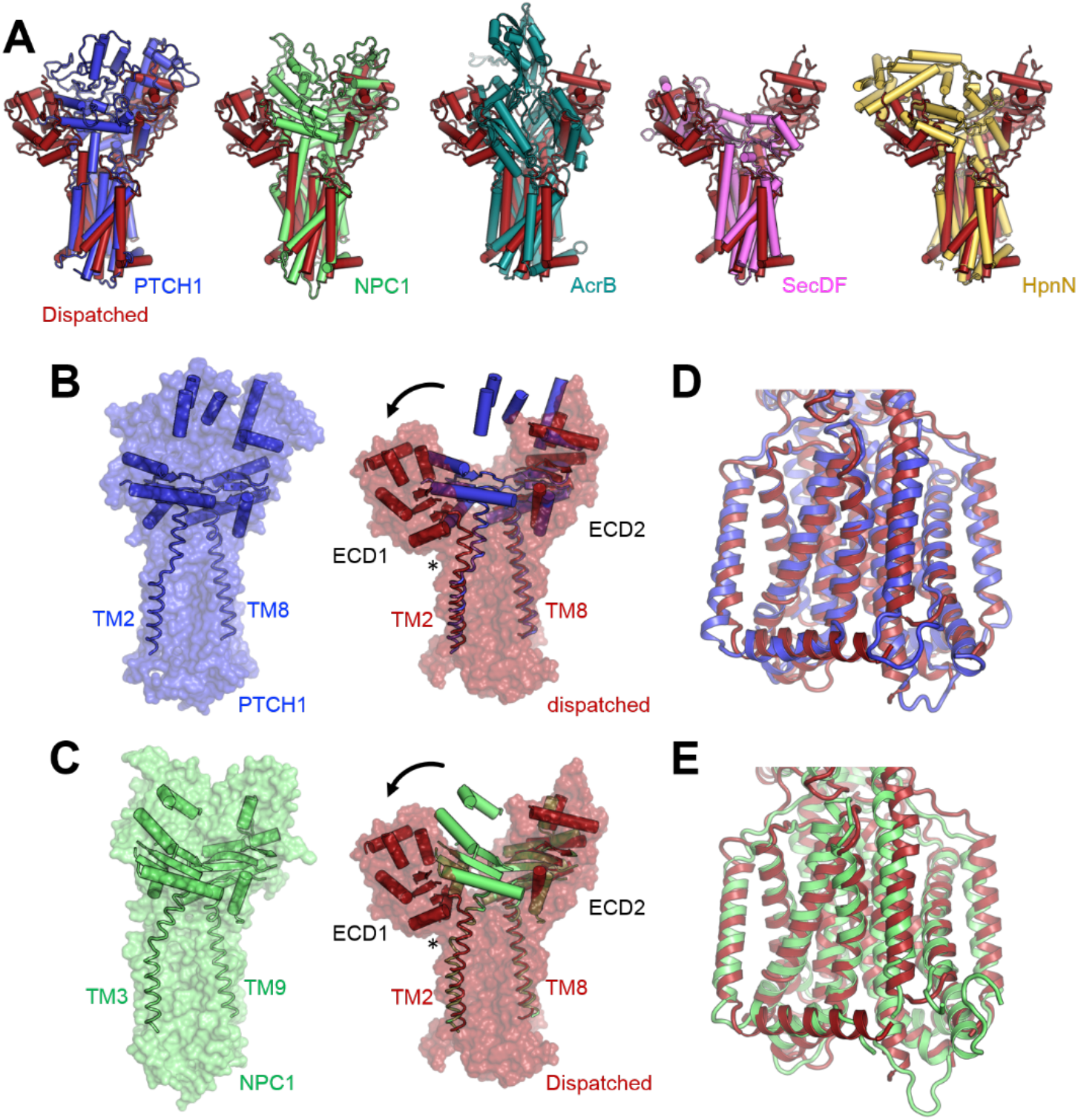
The conformation of Dispatched is unique among the RND proteins with known structures. (**A**) Structural alignment of Dispatched with PTCH1, NPC1, AcrB, SecDF and HpnN reveals a remarkably diverse arrangement of the ectodomains in these proteins. (**B-C**) Comparison to the closest homologues of Dispatched, NPC1 (B) and PTCH1 (C), shows that the ECD1 of Dispatched swings out to create the ECD cavity. The hinge region for this rearrangement appears to be at the boundary between the C-terminus of ECD1 and the N-terminal portion of TM2 of Dispatched (indicated by an asterisk). The major secondary structure elements, α-helices and β-sheets, present in the ectodomains of Dispatched, PTCH1 and NPC1 are shown in cartoon representation. The connecting TM domains 2 and 8 (3 and 9 in NPC1) are shown as ribbons. (**D-E**) Despite the large differences in the relative position of ECD1, the transmembrane domains of Dispatched, PTCH1 and NPC1 are very well aligned.

### Comparisons to RND transporters of known structure reveal a novel open conformation of the Dispatched ectodomain

Comparison of the Dispatched atomic model to those of other RND transporters solved previously by either X-ray crystallography or cryo-EM, including human PTCH1 (pdb id: 6d4h), human NPC1 (pdb id: 5u74), *Escherichia coli* AcrB (pdb id: 2hrt), *Thermus thermophilus* SecDF (pdb id: 3aqp), and *Burkholderia multivorans* HpnN (pdb id: 5khn) revealed a distinct arrangement of the ECD region of Dispatched (Fig. 2A). The closest structural homologues of Dispatched, PTCH1 and NPC1, adopt a closed conformation of the ECD (Fig. 2B-C). The open conformation of Dispatched is established by a splaying motion of the ECD1 relative to ECD2 and the TM domain bundle, at a hinge point residing in the region of the protein at the TM2-ECD1 boundary (Fig. 2C-D). Despite this dramatic difference in the position of the ECD1, the TM domains of Dispatched and PTCH1 or NPC1 can be accurately aligned (Fig. 2D-E).

### Structure of a Dispatched-Hedgehog complex reveals unique protein-based interactions

Given the role of Dispatched in Hh release, we reasoned that the ECD cavity may have a role in binding of the ligand. To test this experimentally, we performed cryo-EM analysis and 3D reconstruction of a complex formed by Dispatched and a recombinant HhN_C85II_ fragment (Fig. 3A, C, Fig. S1B, S5, S6). The modified ligand, lacking a sterol modification and featuring two Ile residues mimicking the palmitoylated C85 residue, was used in analogous fashion to the human ShhN_C24II_ (*24*). In comparison to the transfected UAS-HhN construct or the lipidated recombinant human ShhN fragment, the HhN_C85II_ did not elicit hedgehog pathway activation under cell culture conditions (Fig. 3A) when inoculated in culture medium. It did however show binding to Dispatched in vitro as measured by microscale thermophoresis, with an apparent affinity of 6.2 ± 2.5 μM (Fig. 3A). Our cryo-EM analysis confirmed the binding of the ligand at the interface between ECD1 and ECD2 of Dispatched (Fig. 3B-C). The resolution of the 3D reconstruction was limited to 4.8 Å, but it was sufficient to place and orient a model of HhN into the map (Fig 3C, Fig. S5, Fig. S6A-B). The N-terminus of HhN is close to the base of the Dispatched ECD cavity, and the C-terminus of HhN is adjacent to the SSD (Fig 3D-E), consistent with a possible role of the SSD in cholesterol interaction. Based on the presence of sterol-like density elements in several sites accessible to the SSD in our apo-Dispatched reconstruction (Fig 1B, Fig. S3-4), the C-terminus of the cholesterol-modified HhN may interact with the protein via more than one site. The functional role of the different sterol interaction sites at the protein-lipid interface of Dispatched will require careful future investigation.

**Fig. 3.**
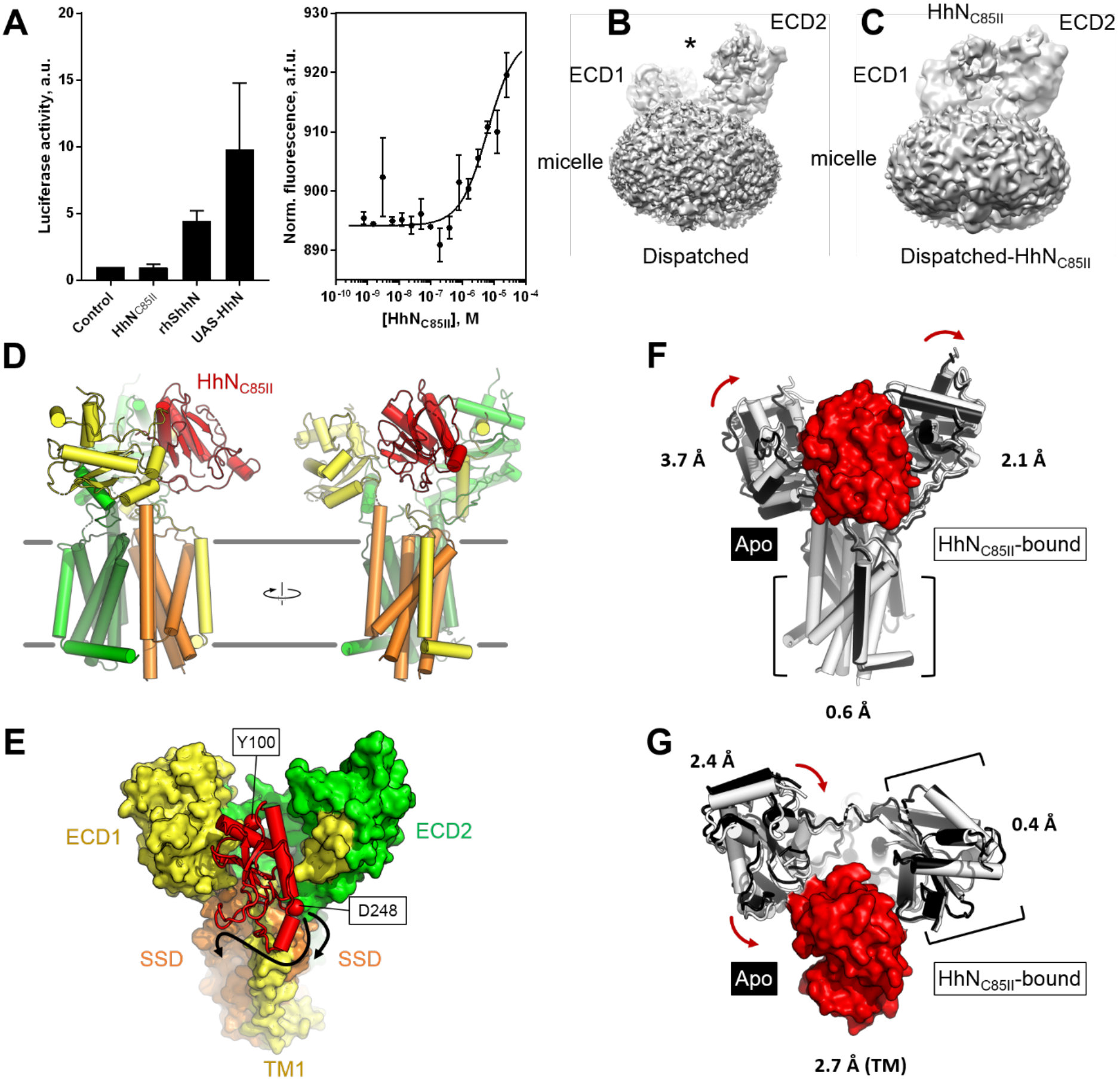
Structure of Dispatched bound to a modified Hedgehog ligand. (**A**) Biochemical activity of HhN_C85II_ ligand, assessed by luciferase luminescence (*left*) and microscale thermophoresis (MST, *right*). The data correspond to values of luciferase activity normalised to the values measured for the “Control” sample (medium without the ligand); n = 4 (for rhShhN n = 2). The affinity of HhN_C85II_ for Dispatched-YFP, determined by MST, was 6.2 ± 2.5 μM (n = 3). (**B-C**) Comparison of the density maps of Dispatched (B) and Dispatched-HhN_C85II_ complex (C); the asterisk (B) indicates the cavity between the ECD1 and 2 regions, occupied by HhN_C85II_ density in the map shown in C. (**D**) Overview of the Dispatched-HhN_C85II_ complex model; colour coding for Dispatched is the same as in Fig. 1C-D; the Hedgehog ligand is coloured red. (**E**) View of the complex from the luminal/extracellular side of the membrane shows the orientation of the HhN_C85II_ ligand. The Cα atoms of N-terminal residue Y100 and the C-terminal residue D248 in the model are shown as red spheres. Arrows indicate possible proximity of the cholesterol-modified C-terminal region of HhN to the SSD region of Dispatched. (**F-G**) Structural alignment of the ligand-free and ligand-bound Dispatched; the complex aligned to apo-Dispatched using the transmembrane domain (F) or using the ECD2 domain (G). The numbers correspond to root mean square deviation (r.m.s.d.) between the atoms of the individual domains. The red arrows indicate the whole domain structural rearrangements that accompany HhN ligand binding.

Only a few residues, H77-H78, R250-253, E296-E298 and N407 of Dispatched ECD1 contribute to the interaction with HhN (Fig. S6). This may explain the relatively low affinity of this interaction and the poor density corresponding to the bound HhN. It is likely that the processed, dually lipidated natural form of HhN engages Dispatched at additional sites within and near the ligand binding cleft. For example, the palmitoylated N-terminus of the protein may be in contact with the surface of ECD2, whereas the C-terminal cholesteryl moiety may interact with membrane-embedded SSD of Dispatched.

We analysed the interfaces that the HhN ligand presents to different binding partners, in the context of our Dispatched-HhN complex (Fig. S7). Structural alignment of the available structures featuring the insect or mammalian Hh ligand bound to its receptors showed that Dispatched interacts with HhN in a unique way (Fig. S7A). Interestingly, two sites in the sequence of the Hedgehog ligands, corresponding to two loop regions in the Drosophila HhN, participated in interactions with all of the compared receptor proteins (Fig. S7B-C). This suggests that these two sites in the morphogen have a critically important role in the Hedgehog pathway, facilitating the interactions between HhN and a wide range of Hedgehog receptor proteins.

## Discussion

The structures of Dispatched alone and in a complex with HhN protein fragment point to exciting new possibilities for the role of Dispatched at the early stage of the Hh signaling and provides the structural basis for understanding the activity of Dispatched (Fig. 4). The conserved arrangement of the core residues previously shown to be important for Dispatched activity (*17*) as well as the presence of multiple sterols at binding sites surrounding the membrane domain of Dispatched point to a striking similarity between the membrane-spanning regions of the eukaryotic RND proteins (Fig. 4, Fig. S3-4). The open conformation of the Dispatched ectodomain capable of accommodating HhN fragment in the cleft between the ECD1 and ECD2 suggests that Dispatched may act directly as a receptor for HhN protein, independently of lipid modification. Although cholesterol-based interactions have been suggested to be critical for ligand release by Dispatched, it is now clear that HhN can engage Dispatched via protein-protein interactions. This interaction may be relevant during the endosomal Dispatched-dependent recycling of HhN (*25*, *26*). Regardless of the precise location where Dispatched acts to facilitate Hh release, the conformation observed in our reconstruction likely represents a pre-release state of the complex.

**Figure. 4.**
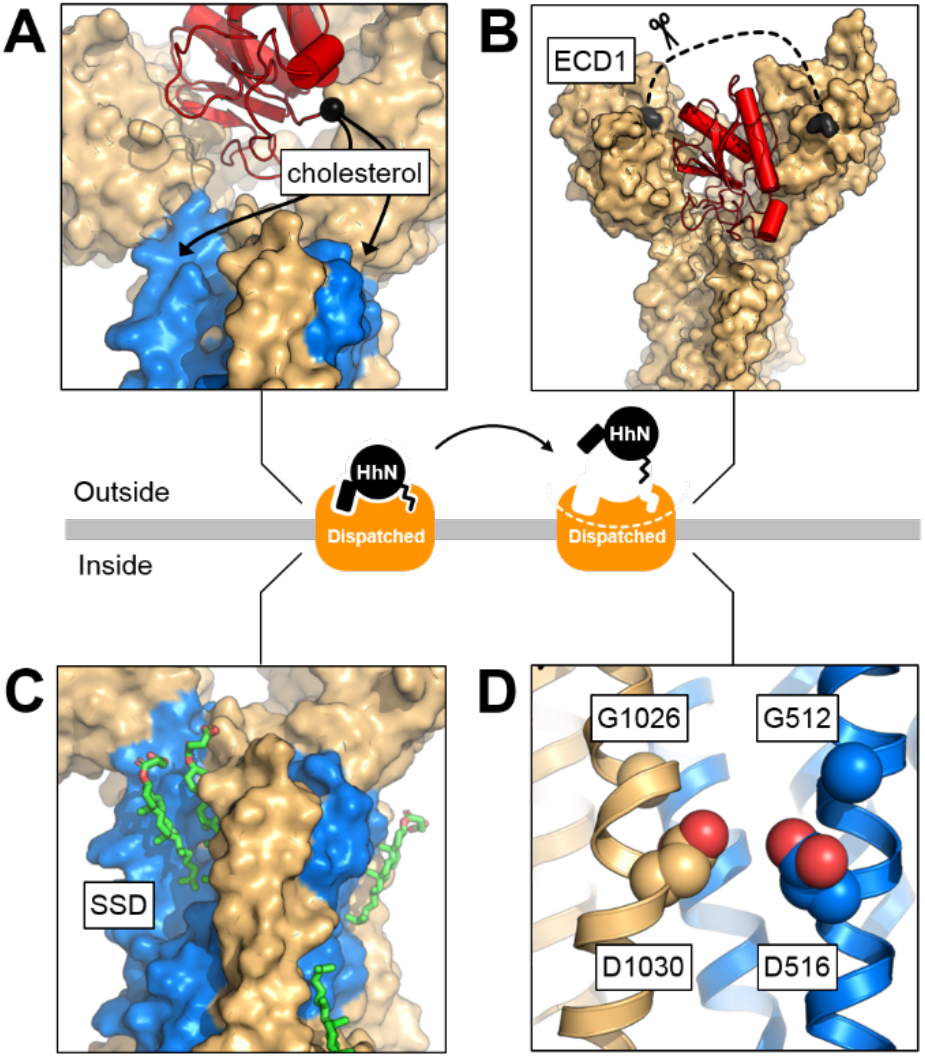
Structural basis of Dispatched activity. The cryo-EM structures of Dispatched alone and in a complex with HhN_C85II_ provide the framework for understanding the structural basis of Dispatched-assisted HhN release. (**A-D**), The key features of Dispatched involved in the activity of Dispatched include: (A) the orientation of the C-terminal region of Hedgehog coincides with the sterol-modification in close proximity to the SSD (blue), (B) the unstructured region of the ECD1 (residues E90-S247; dashed line), which may represent a molecular gate for access or dissociation of the Hedgehog ligand; proteolytic processing of this region has been suggested as a prerequisite for Dispatched activation (*18*), (C) multiple sterols identified the model of apo-Dispatched, (D) the GxxxD motifs in TM4 and TM10, presumed to be involved in transport-like activity of Dispatched (*17*). The release of the Hedgehog ligand is likely dependent on the interplay between these structural elements of Dispatched.

A number of questions pertaining to the structure of Dispatched remain open. For example, our preparations of the protein clearly show that it is monomeric in detergent micelle, yet it has previously been suggested to form oligomers (*18*, *27*). In light of the variety of stable oligomeric states observed recently by cryo-EM for PTCH1, including monomers (*19*, *21*, *24*), dimers (*20*, *22*) and tetramers (*28*), it is possible that Dispatched is capable of oligomerizing in a ligand-dependent or -independent manner. On the other hand, although the structural features and the biological role of Dispatched appear to be unique among the RND transporters, it is also possible that homologous proteins (e.g., PTCH1 and NPC1) may under certain conditions adopt open conformations similar to that of Dispatched.

## Materials and Methods

### Expression and purification of Drosophila Dispatched

The synthetic DNA fragment encoding the D. melanogaster Dispatched (Genewiz; UniProt ID: Q9VNJ5) was cloned into a pACMV vector (*29*) in frame with a C-terminal 3C-YFP-twinStrep tag fusion. The plasmid was transfected into HEK293F cells and a stable clone of the cells constitutively expressing Dispatched was selected by treatment of the cells with G418 according to the established protocol (*29*). Protein expression was performed in 2L Erlenmeyer flasks, with 800 ml culture per flask, at 37°C in Protein Expression Medium (PEM) in the presence of 5% Fetal Calf Serum (FCS), 1x penicillin/streptomycin mixture, 1x Glutamax and 1x Non-essential Amino Acids (NEAA). At cell density of ~2 × 10^6^ cells/ml, the cultures were collected by centrifugation at 500 × g, resuspended in PBS (40 ml per 1 L of culture), frozen and stored at -80°C until purification.

For protein purification, the frozen pellets corresponding to 4L culture of Dispatched-expressing HEK293F cells were thawed, resuspended in 150 ml of buffer A (Tris pH 8, 50 mM, NaCl 150 mM, glycerol 10%), in the presence of 1 mM PMSF and 1 tablet per 50 ml of Roche complete protease inhibitor cocktail, EDTA-free). The resuspended cells were homogenized using a Dounce homogenizer. Following an incubation on a rotating wheel in the presence of 1% DDM and 0.2% cholesteryl hemisuccinate (CHS) at 4°C for 1h, the cell lysates were clarified by ultracentrifugation (rotor Ti45, 35000 rpm, 4°C, 30 min). The supernatants were added to 3 ml of pre-equilibrated CNBr-sepharose resin coupled with purified anti-GFP nanobody (*30*) and incubated for 1 h on a rotating wheel at 4°C. The sepharose resin was collected by gravity flow into a glass Econo-Column (BioRad), washed with 30 column volumes of buffer B (Tris pH 8, 50 mM, NaCl 150 mM) containing DDM/CHS mixture (0.025% DDM / 0.005 % CHS), and with 10 column volumes of buffer B containing 0.1% digitonin. The protein was eluted from the resin using overnight cleavage by 3C protease at 4°C, and the eluted protein-containing solution was subjected to a negative NiNTA purification step (using 1 ml of NiNTA resin pre-equilibrated with buffer B) to remove the 3C protease. The final purified protein sample was isolated using size exclusion chromatography (SEC) using Superose 6 Increase 10/300 GL column equilibrated in buffer B in the presence of 0.1% digitonin (Fig. S1). The selected fractions of the SEC peak corresponding to the purified Dispatched were concentrated to 4 mg/ml using an Amicon Ultra-4 concentrator (100 kDa cut-off, Millipore).

For purification of Dispatched-YFP, the frozen pellets of Dispatched-expressing cells (2L of culture) were thawed and resuspended in 60 mL of Buffer A in presence of 1 mM PMSF and 1 tablet of Roche complete protease inhibitor cocktail (EDTA-free). The resuspended cells were homogenized using a Dounce homogenizer. Following an incubation on a rotating wheel in the presence of 1% DDM and 0.2% cholesteryl hemisuccinate (CHS) at 4°C for 1h, the cell lysates were clarified by centrifugation (rotor A8.24, 19000 rpm, 4°C, 30 min). The supernatant was incubated with 5 mL of equilibrated StrepTactin Superflow resin (IBA Lifesciences). The resin was washed and equilibrated with Buffer A containing 0.02% DDM and 0.004% CHS. Soluble lysate was mixed with the resin and incubated for 30 min, and then loaded on a gravity column. Resin was washed with 3 × 10CV (column volumes) of Buffer A containing 0.02% DDM and 0.004% CHS, and 2 × 5CV of Buffer A containing 0.1% Digitonin. Elution was performed in the same buffer supplemented with 5 mM desthiobiotin (DTB). The protein was further purified using size-exclusion chromatography on a Superose 6 Increase 10/300 GL column equilibrated in buffer B (Tris pH 8, 50 mM, NaCl 150 mM) in the presence of 0.1% digitonin. Selected fractions were concentrated to 1 mg/mL using Amicon Ultra-4 concentrator (100 kDa cut-off, Millipore), and flash-frozen in 10% glycerol.

### Expression and purification of Drosophila hedgehog N-terminal fragment HhN

The synthetic DNA fragment encoding the modified *D. melanogaster* hedgehog N-terminal fragment, referred to as HhN (residues 85-257, modified at the N-terminus by replacing the Cys85 residue with two Ile residues; UniProt ID: Q02936), fused with the N-terminal 6xHis-SUMO tag, was cloned into pET28a plasmid. For expression of the HhN protein, the plasmid was transformed into BL21(DE3) RIPL E.coli cells. The transformed cultures were grown in a shaking incubator at 37°C until OD600 of 0.8, at which point the temperature was switched to 30°C and the expression was induced with 1 mM IPTG for 3h. The induced cells were harvested by centrifugation, frozen and stored at -80°C until the day of experiment.

The frozen E.coli pellets (corresponding to 4 L cultures) were thawed, resuspended in 120 ml buffer C (50 mM Tris, pH 7.5, 200 mM NaCl) containing 25 mM imidazole, disrupted by sonication, and clarified by centrifugation at 18000 g for 30 min at 4°C. The supernatant was added to 2 ml of NiNTA resin, incubated at 4°C with rotation for 1 h. The NiNTA resin was collected into a gravity column, washed with 30 column volumes of buffer C containing 25 mM imidazole and 10 column volumes of buffer C containing 50 mM imidazole. The protein was eluted using buffer C containing 200 mM imidazole. The eluted fractions were pooled, diluted 8-fold to reduce the concentration of imidazole and incubated with 250 μg of purified SUMO protease Ulp1 overnight at 4°C. The mixture was passed though 2 ml of immobilized NiNTA resin pre-equilibrated with buffer C containing 25 mM imidazole to remove the uncleaved HhN, the cleaved off SUMO tag and the Ulp1. The flow-through was concentrated to 1 ml and applied to a Superdex 200 Increase 10/300 GL column (Fig. S1). The fractions corresponding to HhN were pooled, supplemented with 10% glycerol, flash-frozen in liquid nitrogen and stored at -80°C until the day of experiment.

### Microscale thermophoresis

For microscale thermophoresis (MST) analysis, the dilution series of purified HhN_C85II_ were prepared in buffer A containing 0.1% digitonin (16 concentration points, ranging from 50 μM to 1.52 nM). Dispatched-YFP was mixed with each solution at a 1:1 ratio (final Dispatched-YFP concentration: 20 nM), and the mixtures were incubated for about 20 min at room temperature. Monolith NT 1.115 Premium Capillaries (NanoTemper) were filled with each reaction mixture, loaded on a cartridge into the Monolith NT 1.115 (NanoTemper) and MST analysis was performed using the following parameters: MST-power medium, excitation-power 100%, excitation type “Blue”, temperature 22 °C. The PALMIST software (*31*) was used for data analysis and model fitting, using weighted fitting for the three sets of data, and with cold values measured before laser firing and hot values after thermophoresis. Analysis results were exported and plotted with Prism.

### Cryo-EM sample preparation and data collection

The concentrated sample of Dispatched (4 mg/ml) was kept on ice and was immediately used for cryo-EM grid preparation. The Dispatched-HhN sample was prepared by mixing the concentrated Dispatched (4 mg/ml) with a 2-fold molar excess of the concentrated HhN (desalted into the buffer B containing 0.1% digitonin), and incubating the mixture for ~30 min on ice prior to cryo-EM grid freezing. For cryo-EM sample preparation, UltrAuFoil 1.2/1.3 Au 300-mesh grids were glow discharged using a PELCO easiGlow (Ted Pella) plasma cleaning system for 30s at 25 mAmp in air. The protein samples (3.5 μl) were deposited on the surface of the grid immediately before blotting for 3 s on a Vitrobot Mark IV (FEI) set to operation at 4°C with 100% humidity and with a blotting force of 2, and plunging the grid into liquid ethane. The grids were stored in liquid nitrogen until the day of data collection.

For Dispatched, a cryo-EM dataset was collected at EMBL Heidelberg, using Titan Krios microscope equipped with a Gatan K2 Summit detector and an energy filter. A total of 9216 micrographs were automatically collected in counting mode using SerialEM, at a magnification of 165000x (resulting in the micrograph pixel size of 0.81 Å/pix), with a total exposure of 8s over 40 frames and a total dose of 50.42 e/Å2.

The a cryo-EM dataset for Dispatched-HhN was collected at NeCEN, Leiden, using a Cs-corrected Titan Krios microscope equipped with a Falcon 3 camera. A total of 5609 movies were collected in counting mode using EPU, at a nominal magnification of 96kx (resulting in the micrograph pixel size of 0.88 Å/pix), with a total exposure of 60s over 49 frames and a total dose of 51.5 e/Å2.

### Cryo-EM data analysis

Drift correction and alignment of the cryo-EM movies was performed using MotionCor2 (*32*), and the defocus values of the aligned micrographs were estimated using Gctf (*33*). Only micrographs with an estimated resolution below 4 Å were kept for further processing using relion-2.1 and relion-3.0 (*34*, *35*). After manual picking and 2D classification using a small subset of data (35554 particles), about 2.2M particles were autopicked from all selected micrographs using the best 2D classes as references. Several rounds of 2D classification of 2-fold downsampled particles reduced the dataset to 531k particles. For the first round of 3D classification, we used an initial model generated in relion-2.1 using data collected with the Jeol JEM2200 at PSI. The best resulting 3D class was used for downstream image processing steps. Following additional rounds of 3D classification, the particles belonging to the best 3D class could be refined to 3.9 Å (FSC cutoff 0.143). Particles were then reextracted and rescaled to the original pixel size, and refined with a mask to 3.2 Å. Bayesian particle polishing and CTF refinement in relion-3.0 resulted in the postprocessed map of Dispatched at 3.15 Å resolution (using b factor of -50 for map sharpening). The complete data processing work-flow is shown in Fig. S2.

For Dispatched-HhN dataset, the initial steps were similar to those described above. After several rounds of 2D classification in relion-3.0, a dataset of 162907 particles was selected. One rounds of 3D classification with 2 classes was performed, and the best 3D class (98623 particles) could be refined to 7.04 Å resolution. This class was subjected to a 3D classification with a single class (40 iterations, T20, E7). The particles in this class were imported into cisTEM(*36*), local searches were performed using an initial resolution cut-off of 15 Å, followed by the same procedure with a 10 Å cut-off. A mask excluding the micelle but covering Dispatched and the bound HhN density was used (a soft edge of 10 Å was applied during processing in cisTEM). The density outside the mask was low pass filtered to 30 Å, and the weight outside the mask was set to 0.3. The refinement converged after several iterations to an overall resolution of 4.3 Å (FSC cutoff 0.143); B-factor sharpening resulted in a final map at 4.76 Å resolution.

### Model building and validation

Model building was performed in Coot (*37*) using the postprocessed map of Dispatched. Building of the Dispatched ECD1 and ECD2 was assisted by homology models generated using SwissModel server using PTCH1 (PDB ID: 6d4h) and NPC1 (PDB ID: 5u74) as templates. The density elements corresponding to bound detergents/lipids were interpreted as molecules of CHS. The atomic model of Dispatched was refined using phenix.real_space_refine (*38*) implemented in Phenix (*38*). For model validation, the atom coordinates in the refined model were randomly displaced by a maximum of 0.5 Å using the PDB tools in Phenix. The derived model was subjected to real space refinement using one of the refined half maps (half-map1). Map vs model FSC comparison was made for the model against the corresponding half-map1 used in the refinement job, and for the same model versus the half-map2 (not used during refinement) (*39*). The geometry of the Dispatched model was validated using MolProbity (*40*). For modeling of the Dispatched-HhN complex, a D. melanogaster hedgehog model (pdb id: 2ibg) and our model of Dispatched were manually docked into the density map in UCSF Chimera and jiggle fit as a rigid body in coot. Figures featuring the models and density maps were prepared using PyMol (*41*) and UCSF Chimera (*42*).

### Dual luciferase activity assay

Clone 8 cells were grown on 10 cm plates in Shields and Sang M3 insect Medium (Sigma #S8398) supplemented with 0.5 mg/mL insulin (Sigma #I6634), 2% FBS (JRH #12103-78P), 2.5% fly extract (Drosophila Genomics Resource Center, Indiana) and penicillin/streptomycin (Gibco #15070-063). A day before transfection, cells were seeded into a 24-well plate at a density of 0.4 × 10^6^ cells/well. On the day of transfection, cells were transfected using the Effectene kit (Qiagen), following the standard protocol. A DNA mastermix was prepared for each assay, with a 5:1 ratio of ptc Firefly luciferase: tub Renilla luciferase constructs. Transfected luciferase or hedgehog plasmids were generated as described previously(*43*); the UAS-HhN construct is constitutively expressed in the transfected cells through regulation of the Upstream Activating Sequence, UAS, by Gal4. For experiments involving purified ligands, purified HhN_C85II_ or recombinant human sonic hedgehog, rhShhN (R&D Systems #8908-SH/CF), was sterile-filtered and added to the medium during transfection to a final concentration of 200 nM. Cells were lysed using the Passive Lysis Buffer from the Dual-Luciferase kit (Promega) after 24h in the case of stimulation with purified ligand, or 48h in the case of ligand transfection. Cell lysates were used immediately upon preparation or stored at -80°C. Luciferase activity was measured according to the Dual-Luciferase kit protocol using PHERAstar FSX microplate reader (BMG Labtech).

## Supplementary Materials

Fig. S1. Purification of Dispatched and the modified hedgehog ligand.

Fig. S2. Cryo-EM and single particle analysis of purified Dispatched.

Fig. S3. Features of the high resolution cryo-EM map and model of Dispatched.

Fig. S4. Three of the observed sterol densities in the 3D reconstruction of Dispatched are conserved in mammalian RND transporters.

Fig. S5. Cryo-EM and single particle analysis of Dispatched-HhN_C85II_ complex.

Fig. S6. Details of the interaction between HhN_C85II_ and Dispatched.

Fig. S7. Structural alignment of the hedgehog complexes reveals a unique Dispatched-HhN interaction.

Table S1. Cryo-EM data collection, single particle analysis and model building statistics.

Movie S1. Cryo-EM density map of Dispatched.

## General

We thank the Electron Microscopy Facility at PSI, Villigen (Elisabeth Mueller-Gubler, Takashi Ishikawa) for support. We thank the EMBL Heidelberg Cryo-Electron Microscopy Service Platform (Felix Weis) for the support and expertise in high resolution cryo-EM data collection. We thank the staff at the NeCEN (Christoph Diebolder) for the support and expertise in high resolution cryo-EM data collection.

## Author contributions

F.C. designed and performed the experiments, analysed the data, wrote the manuscript.

C.Q. performed the experiments.

J.F. performed the experiments, analysed the data.

G.H. designed the experiments, analysed the data.

K.B. designed the experiment, analysed the data.

V.M.K. designed and performed the experiments, analysed the data, wrote the manuscript.

## Data and materials availability

Coordinates and cryo-EM maps will be deposited in the Protein Data Bank and Electron Microscopy Data Bank. All other data is available in the manuscript or the supplementary materials. Correspondence and material requests should be addressed to.

