## Supplementary Materials for "Cryo-EM structure of the Hedgehog release protein Dispatched"

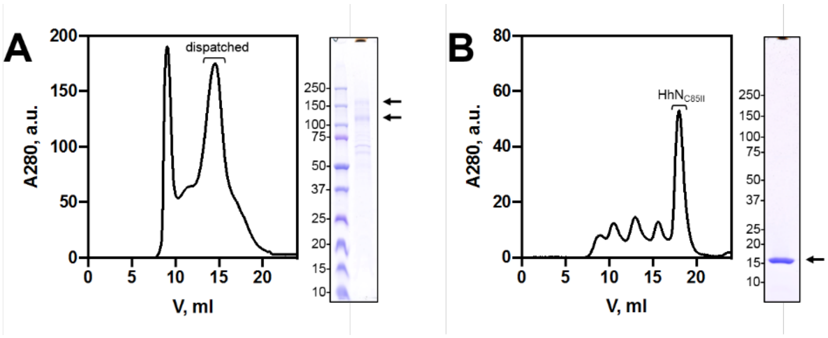


**Fig. S1. Purification of Dispatched and the modified hedgehog ligand**. (**A**) Size exclusion chromatography (SEC) and SDS PAGE of Dispatched. For SEC, a Superose 6 10/300 GL Increase column was used. Protein purity was analysed using a 4-20% gradient SDS-PAGE gel. The arrows indicate the bands corresponding to Dispatched; the protein (loaded upon addition of the loading buffer, without heating) migrates as two bands in SDS PAGE gels. (**B**) Same as in (A) for the recombinant $HhN_{C85II}$ purified using Superdex 200 10/300 GL Increase and analysed using a 4-20% gradient SDS-PAGE gel. The arrow indicates the purified $HhN_{C85II}$.

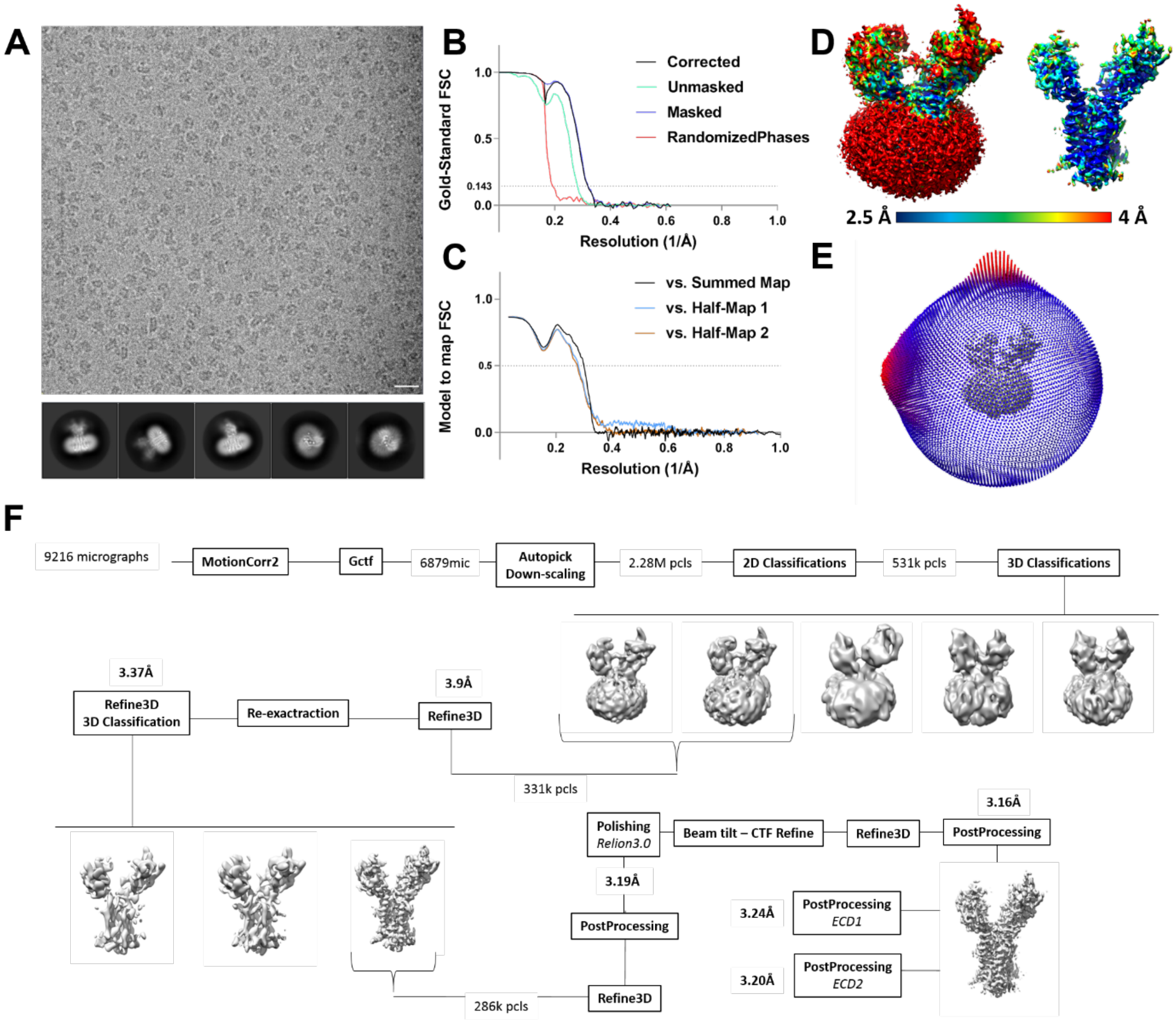


**Fig. S2. Cryo-EM and single particle analysis of purified Dispatched.** (**A**) A representative motion-corrected micrograph of Dispatched (top; scale bar corresponds to 200Å) and a selection of the best 2D classes (bottom; box edge corresponds to 246 Å). (**B**) Fourier shell correlation (FSC) plot for the 3D reconstruction of Dispatched, refined and postprocessed to a resolution of 3.16 Å (FSC 0.143). (**C**) Model to map FSC plot for the Dispatched model, calculated as described in "Methods". (**D**) Local resolution estimate for the processed map by REFMAC (top) and angular distribution of the particles used in refinement (bottom). (**E**) Overview of the image processing workflow.

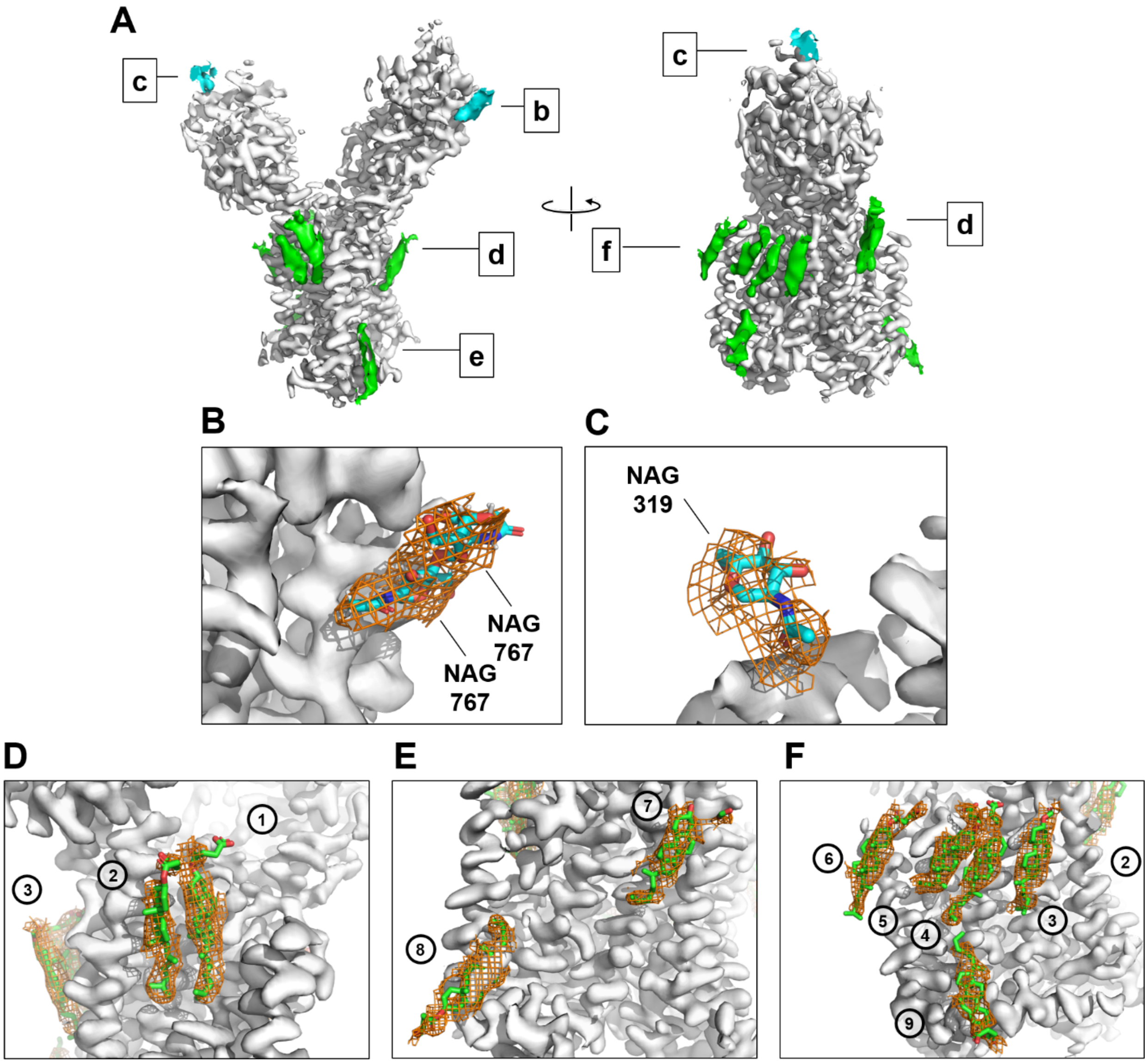


**Fig. S3. Features of the high resolution cryo-EM map and model of Dispatched.** (**A**) Overview of the density map. Dispatched is depicted as a surface display (grey), meshes indicate the sharpened map at 5σ (3σ in panel C). Cyan and green densities correspond to glycosylation sites and bound sterol molecules, respectively. (**B-E**) Zoomed in views of selected features of the density map, including the densities corresponding to a well-resolved glycosylation site (B-C; NAG – N-acetylglucosamine) and the sterols at the protein-lipid interface (D-F; sterol positions are indicated by circles); densities for nine sterol molecules were modeled in the vicinity of the Dispatched membrane domain.

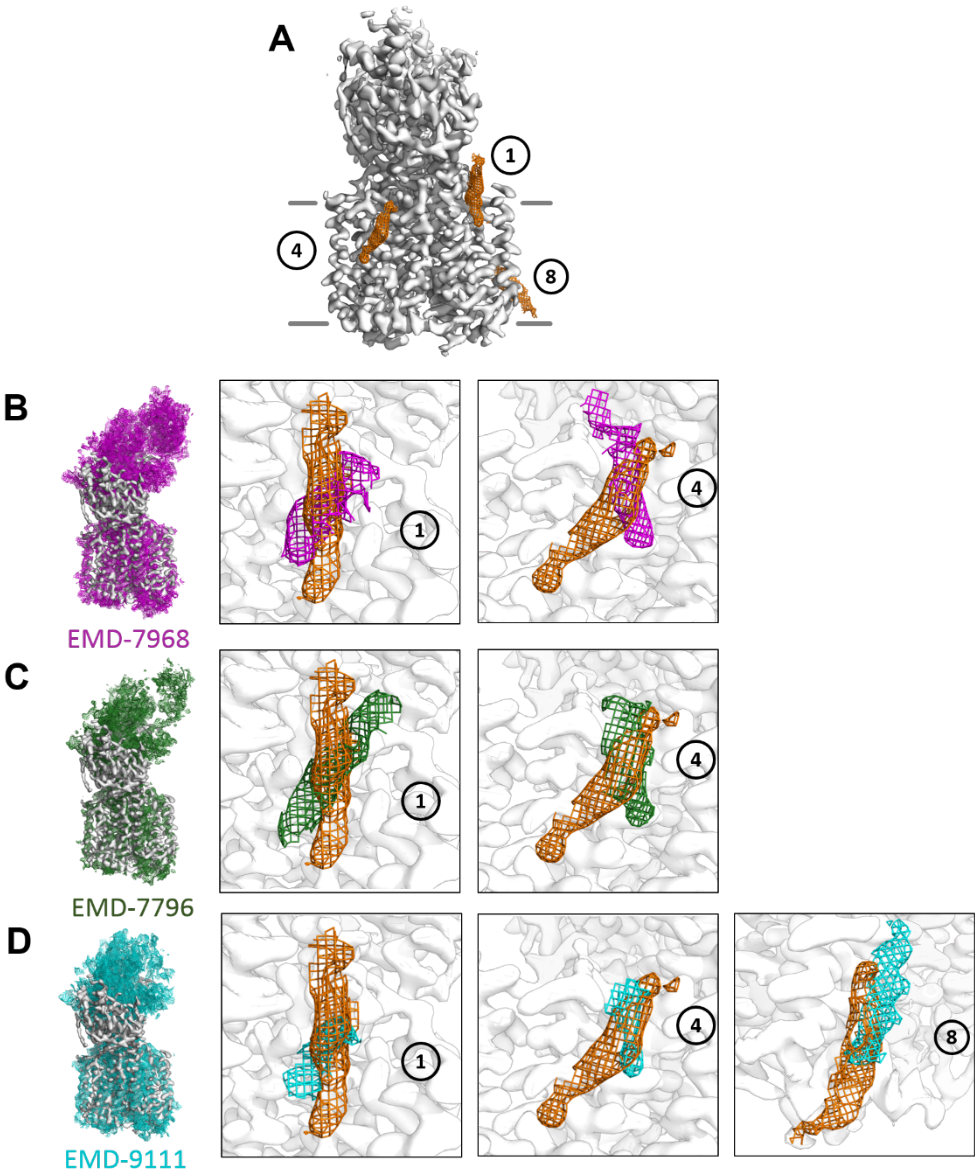


**Fig. S4. Three of the observed sterol densities in the 3D reconstruction of Dispatched are conserved in mammalian RND transporters.** (**A**) Overview of the Dispatched map (grey) and sterol densities (orange) at 5σ. The densities of the matching sterol molecules were identified by inspection of the density maps for PTCH1: PDB ID 6d4j (EMD-7796, green), PDB ID 6dmy (EMD-7968, purple), PDB ID 6mg8 (EMD-9111, blue). The density maps were aligned to the sharpened Dispatched map using the conserved transmembrane domain region. (**B-D**) Zoomed in views of the conserved sterol sites; the labels correspond to those shown in Fig. S3C-E.

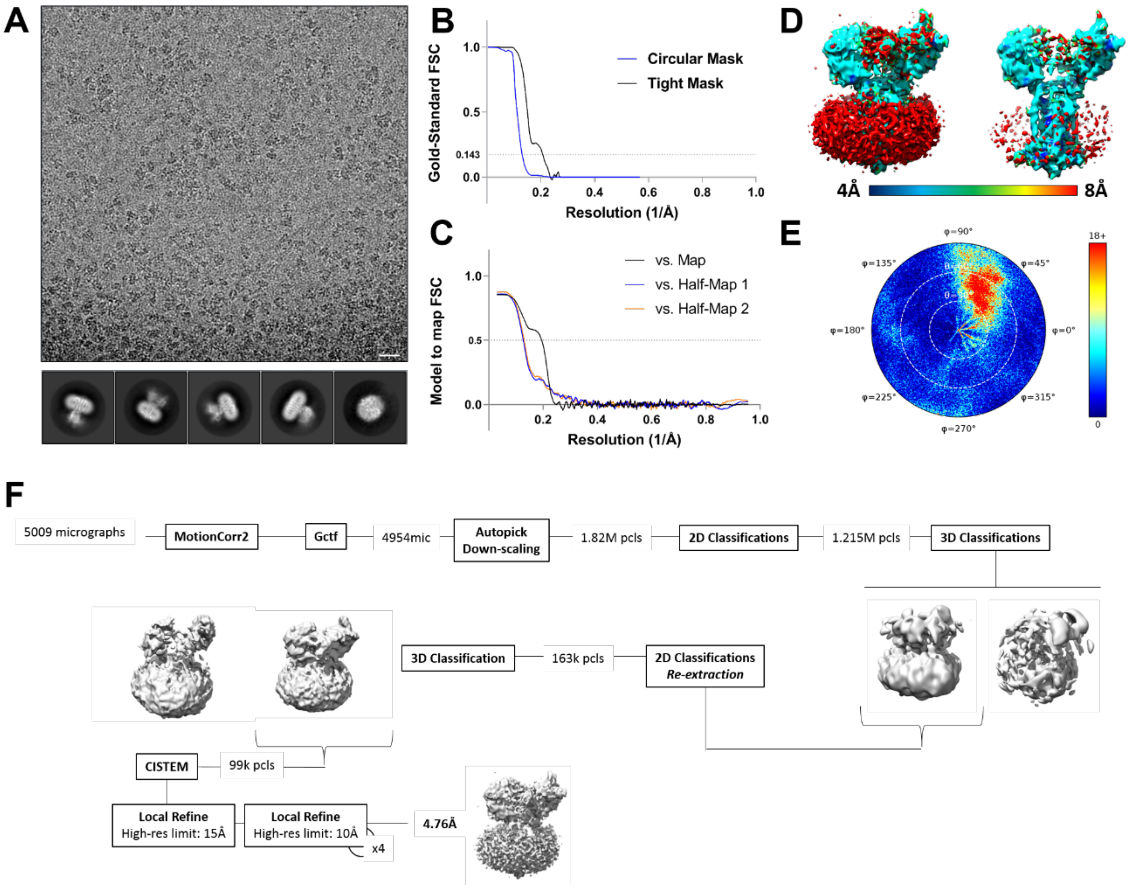


**Fig. S5. Cryo-EM and single particle analysis of Dispatched-$\text{HhN}_{\text{C85II}}$ complex.** (**A**) A representative motion-corrected micrograph of Dispatched-$\text{HhN}_{\text{C85II}}$ complex (scale bar corresponds to 200 Å) and a selection of 2D classes (box edge corresponds to 267 Å). (**B**) Fourier shell correlation (FSC) plot for the refined map of Dispatched (4.76 Å resolution, FSC 0.143). (**C**) Model to map FSC plot of the built models to the half-maps generated by cisTEM. (**D**) Local resolution estimates for the refined 3D reconstruction (top) and angular distribution of the particle dataset (bottom). E**E**) Overview of the image processing workflow. Initial steps of image processing were performed in Relion; 3D refinement of the final particle selection was performed in cisTEM.

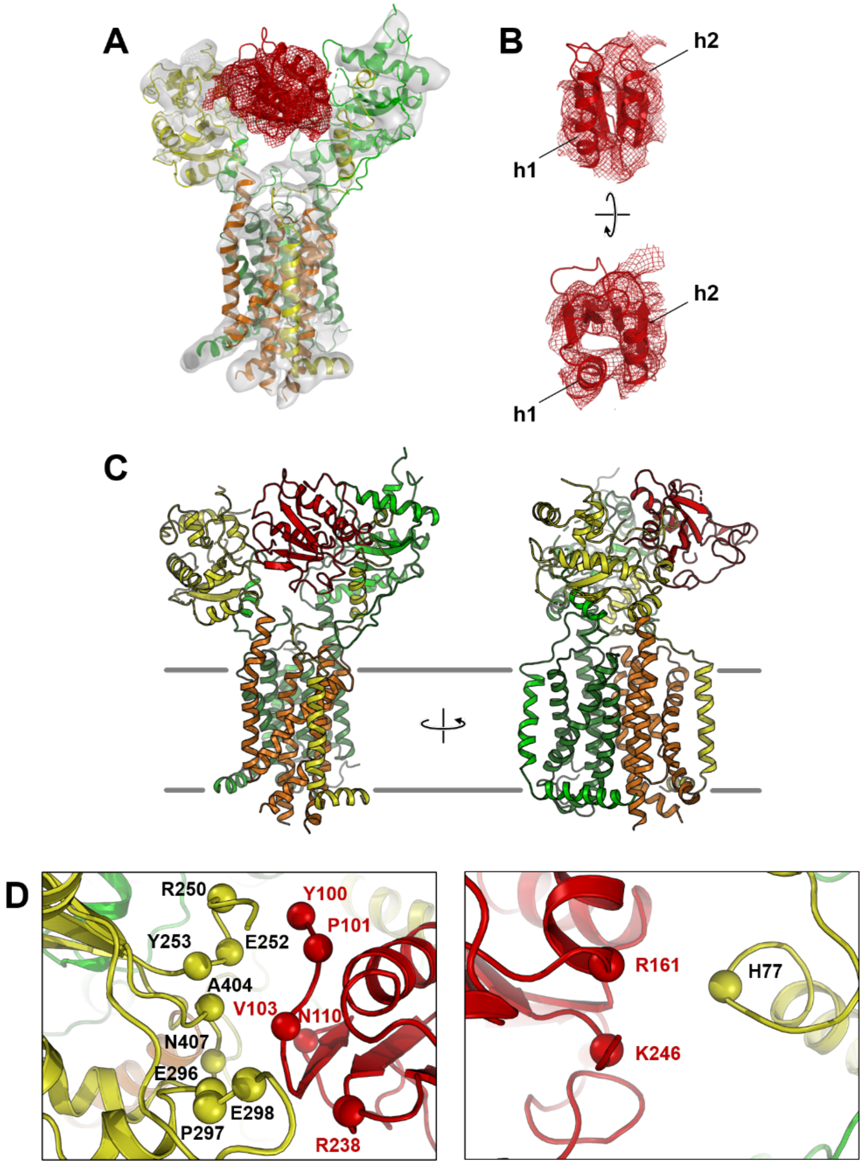


**Fig. S6. Details of the interaction between HhN$_{C85II}$ and Dispatched.** (**A**) A cartoon representation of the models of Dispatched (same colour code as in Fig. 1) and HhN$_{C85II}$ (red), fitted in the density. Density is displayed at 10σ (grey surface) and 5σ (red mesh). (**B**) The model of HhN (PDB ID: 2ibg) modelled into the density map using HhN α-helices for reference. (**C**) Overview of the Dispatched-HhN$_{C85II}$ model; the colors of Dispatched correspond to the colour coding in Fig. 1, Hedgehog is displayed in red. (**D**) Dispatched-HhN interaction interface. Residues involved in the interaction (selected in each chain within 4 Å of the neighbouring molecule) are shown as spheres.

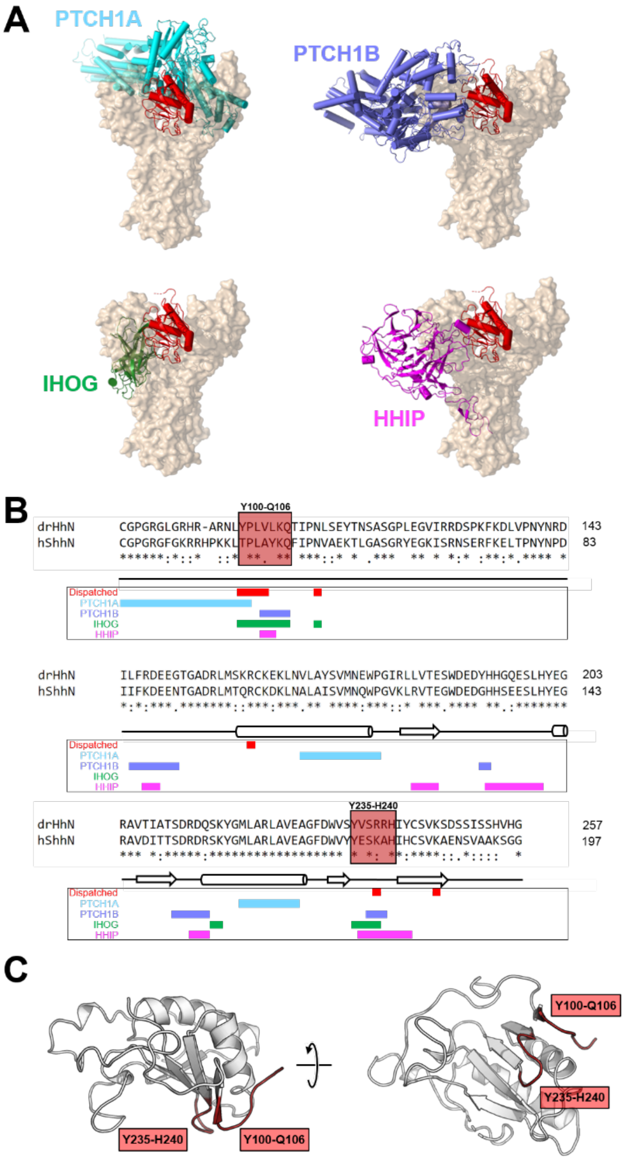


**Fig. S7. Structural alignment of the hedgehog complexes reveals a unique Dispatched-HhN interaction.** (**A**) Alignment of Hh-binding proteins in complex with hedgehog ligands: PTCH1A and PTCH1B (light and dark blue, PDB ID: 6e1h), HHIP (magenta, PDB ID: 3ho5) and Ihog (green, PDB ID: 2ibg), compared to Dispatched (orange). (**B**) Sequence alignment of the human sonic hedgehog N-terminal fragment (ShhN) and drosophila hedgehog (HhN). The secondary structure of distinct portions of the proteins are indicated. The interaction sites of the hedgehog ligands with the respective receptors are coloured as in a. Two clusters of residues participating in the known Hedgehog-binding protein interactions are highlighted with a red box. (**C**) The interaction interface clusters (red cartoon) mapped on the *Drosophila* Hedgehog structure.

| Data collection | | |
|---|---|---|
| **Protein sample** | **Dispatched** | **Dispatched-HhN** |
| **Instrument** | K2 | Falcon III |
| **Magnification** | 165kx | 96kx |
| **Voltage (kV)** | 300 | 300 |
| **Electron exposure ($e^-$/Å$^2$)** | 50 | 51.5 |
| **Defocus range (μm)** | -0.8 -2.4 | -0.7 -2.5 |
| **Pixel size (Å)** | 0.81 | 0.88 |
| **Resolution (Å; FSC 0.143)** | 3.16 | 4.76 |
| **Number of particles** | 286136 | 98623 |
| **Model refinement** | | |
| **Model resolution (FSC 0.5)** | 3.34 | 5.0 |
| **Map sharpening b-factor (Å)** | -50 | -50 |
| **Map CC** | 0.774 | 0.734 |
| **Model composition** | | |
| **Protein residues/ligands** | 842 | 842/140 |
| **B factor (Å$^2$)** | 60.45 | 239.6 |
| **Bond length RMSD (Å)** | 0.010 | 0.007 |
| **Bond angle RMSD (°)** | 1.266 | 1.032 |
| **Validation** | | |
| **MolProbity score** | 1.62 | 1.99 |
| **Clash score** | 4.57 | 9.71 |
| **Rotamer outliers (%)** | 0 | 0.7 |
| **Ramachandran plot** | | |
| **Favored (%)** | 94.3 | 92.26 |
| **Allowed (%)** | 5.7 | 6.38 |
| **Disallowed (%)** | 0 | 0 |

**Table S1.** Cryo-EM data collection, single particle analysis and model building statistics.

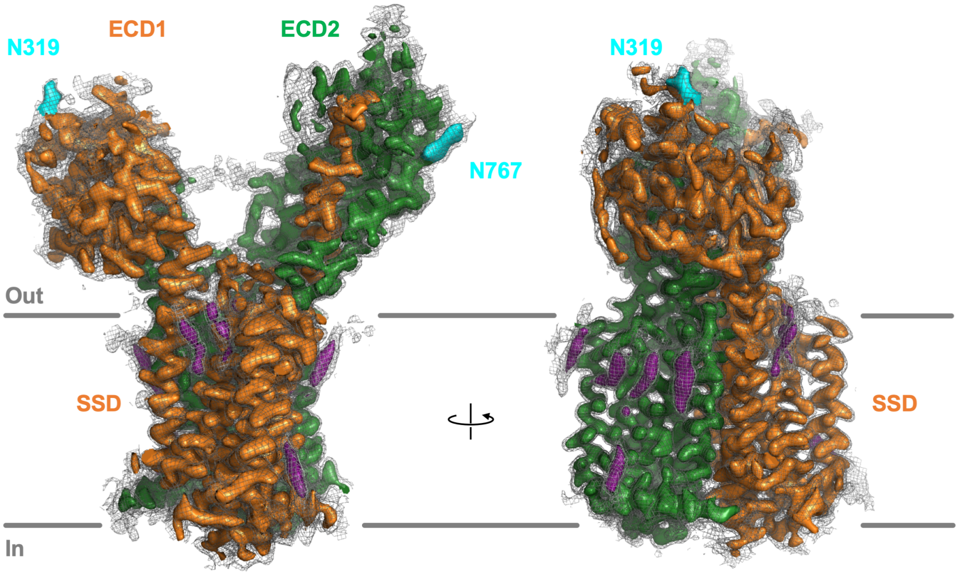


**Movie S1. Cryo-EM density map of Dispatched.** A movie displaying the sharpened density map (sharpened in relion using a b factor of -50 Å$^2$), displayed at 10σ (NAG319 displayed at 5σ). The density elements corresponding to the bound sterols are coloured magenta. The density corresponding to the glycosylation sites are coloured cyan. Grey mesh covering the density map is contoured at level 4.5σ.
